# PloverDB: A high-performance platform for serving biomedical knowledge graphs as standards-compliant web APIs

**DOI:** 10.1101/2025.03.09.642156

**Authors:** Amy K. Glen, Eric W. Deutsch, Stephen A. Ramsey

## Abstract

Knowledge graphs are increasingly being used to integrate heterogeneous biomedical knowledge and data. General-purpose graph database management systems such as Neo4j are often used to host and search knowledge graphs, but such tools come with overhead and leave biomedical-specific standards compliance and reasoning to the user. Interoperability across biomedical knowledge bases and reasoning systems necessitates the use of standards such as those adopted by the Biomedical Data Translator consortium. We present PloverDB, a comprehensive software platform for hosting and efficiently serving biomedical knowledge graphs as standards-compliant web application programming interfaces. In addition to fundamental back-end knowledge reasoning tasks, PloverDB automatically handles entity resolution, exposure of standardized metadata and test data, and multiplexing of knowledge graphs, all in a single platform designed specifically for efficient query answering and ease of deployment. PloverDB increases data accessibility and utility by allowing data providers to quickly serve their biomedical knowledge graphs as standards-compliant web services. **Availability and Implementation:** PloverDB’s source code and technical documentation are publicly available under an MIT License at github:RTXteam/PloverDB, archived on Zenodo at doi:10.5281/zenodo.15454600.

## 1 Introduction

Knowledge graphs (KGs) have emerged as useful abstractions for integrating heterogeneous biomedical data, making them convenient substrates for data mining and other computational reasoning techniques. The way in which such knowledge graphs are exposed for use varies; some graphs are only available for flat-file download, while others are exposed via a web application programming interface (API) or user interface, often backed by a general-purpose graph database management system such as Neo4j (github:neo4j/neo4j). Naturally, a knowledge graph’s ultimate utility is heavily influenced by the convenience, speed, and reliability of its mode of access.

The National Center for Advancing Translational Sciences (NCATS) Biomedical Data Translator [2] (abbreviated as “Translator”) is a distributed knowledge graph-based computing system that integrates and reasons across disparate biomedical data to answer questions like *“What drugs might be repurposed to treat Fanconi anemia?”*. The Translator Consortium has defined data standards for biomedical knowledge graphs, including 1) the *Biolink Model*, a standard schema and semantic layer for biomedical KGs [8], and 2) *TRAPI*, a standard JavaScript Object Notation (JSON) web API format for biomedical KGs (github:NCATSTranslator/ReasonerAPI).

Within Translator, knowledge graphs are exposed via TRAPI-compliant web APIs that can answer single-edge or “one-hop” pattern-matching queries, such as *Acetaminophen—interacts_with* → *Protein?* (see Sec. 2.1.1), and in doing so, perform certain fundamental reasoning tasks like transitive chaining of concept subclass relationships (see Sec. 2.1.2). Such KG APIs can then be used by reasoning systems, such as Translator reasoning agents (e.g., [1], [3], and github:ranking-agent/aragorn) or large language model-based chain-of-thought reasoning systems, to answer larger queries.

As Translator’s requirements for interoperability and graph-based reasoning illustrate, transitioning from a flat-file version of a knowledge graph to a deployed, standards-compliant web API is not a trivial task. The KG owner must 1) canonicalize their graph (i.e., identify and unify semantically identical concept nodes); 2) import it into some sort of database platform (such as Neo4j [6, 9]); 3) convert incoming TRAPI queries into queries runnable on the back-end database system; 4) encode various semantic and other fundamental reasoning tasks such as transitive chaining of subclass relationships; 5) ensure that answers are transformed back into TRAPI format; 6) compute a meta knowledge graph and test triples; and 7) deploy the entire service as a performant web API, even in the face of heavy concurrent load.

Previous work on web-accessible hosting frameworks for biomedical knowledge graphs includes Plater (github: TranslatorSRI/Plater), a tool for exposing a Neo4j-hosted Biolink-compliant KG as a TRAPI web API, which can be com-bined with Babel (github:TranslatorSRI/Babel) and ORION (github:RobokopU24/ORION, part of the ROBOKOP [6] software stack) to achieve entity resolution and KG formation/import, respectively. Another tool, BioThings Software Development Kit [5], can dynamically create a Biolink-compliant API for a KG according to a custom data parser, which can then be exposed as a TRAPI API by BioThings Explorer [1].

Despite previous efforts, there is a need for a platform that allows data owners to easily deploy Biolink-compliant flat-file knowledge graphs as TRAPI web APIs that 1) is *comprehensive* in terms of back-end reasoning (e.g., entity resolution, concept subclass chaining) and provided artifacts/endpoints (e.g., meta KG, test triples), 2) is *highly-performant*, and can be *independently* deployed with *minimal effort* upon KG content or knowledge representation changes. To fill this gap, we created PloverDB, an all-in-one platform for efficiently hosting and serving Biolink-compliant KGs as TRAPI web APIs. We describe PloverDB’s implementation and usage in the following sections.

## 2 Implementation

PloverDB is a fully in-memory Python-based platform for hosting and serving Biolink-compliant knowledge graphs as TRAPI web APIs, designed specifically for improved query speed and ease of deployment. Its design prioritizes ease of use and minimal configuration, enabling rapid deployment of KGs with minimal overhead. PloverDB was initially created in 2021 to replace Neo4j as the back-end database platform for the RTX-KG2 knowledge graph [9]^1^, but was generalized in 2024 to 1) act as a standalone, fully TRAPI-compliant web service and 2) be compatible with any Biolink-compliant knowledge graph.

PloverDB is Dockerized for convenient deployment, as depicted in Figure 2.1. During its Docker image build, PloverDB 1) downloads the Biolink-compliant KG nodes and edges files—which may be in either TSV or JSON Lines Knowledge Graph Exchange (KGX) format (github:biolink/kgx)—from user-specified public URLs provided in a JSON configuration file; 2) loads the graph into memory; 3) builds its core data structure and other indexes, utilizing the Biolink Model and Translator Node Normalizer (github:TranslatorSRI/NodeNormalization) service as applicable; optionally canonicalizes the graph; and 5) saves indexes to disk within the image. When a Docker container is then run from that image, PloverDB loads its core data structure and indexes into memory and starts a uWSGI/Flask server that exposes the graph via a TRAPI-compliant API, providing various endpoints (only standardized endpoints are depicted in Figure 2.1).

PloverDB’s core data structure is essentially a nested adjacency dictionary in which nodes are mapped to their neighbors, organized by the categories of those neighbors and the types of edges that connect them to those neighbors. PloverDB does not use any intermediary database platforms such as Neo4j or SQLite, and instead stores the graph in Python data structures (dictionaries and sets). At runtime, multiple uWSGI workers (processes) share these read-only data structures to handle concurrent requests.

PloverDB is configured for a given knowledge graph via a single JSON config file, where users provide public URLs from which their KG files should be downloaded, can specify how node/edge properties should be represented in TRAPI attributes, and can control other options like graph canonicalization (see Sec. 3).

**Figure 2.1:**
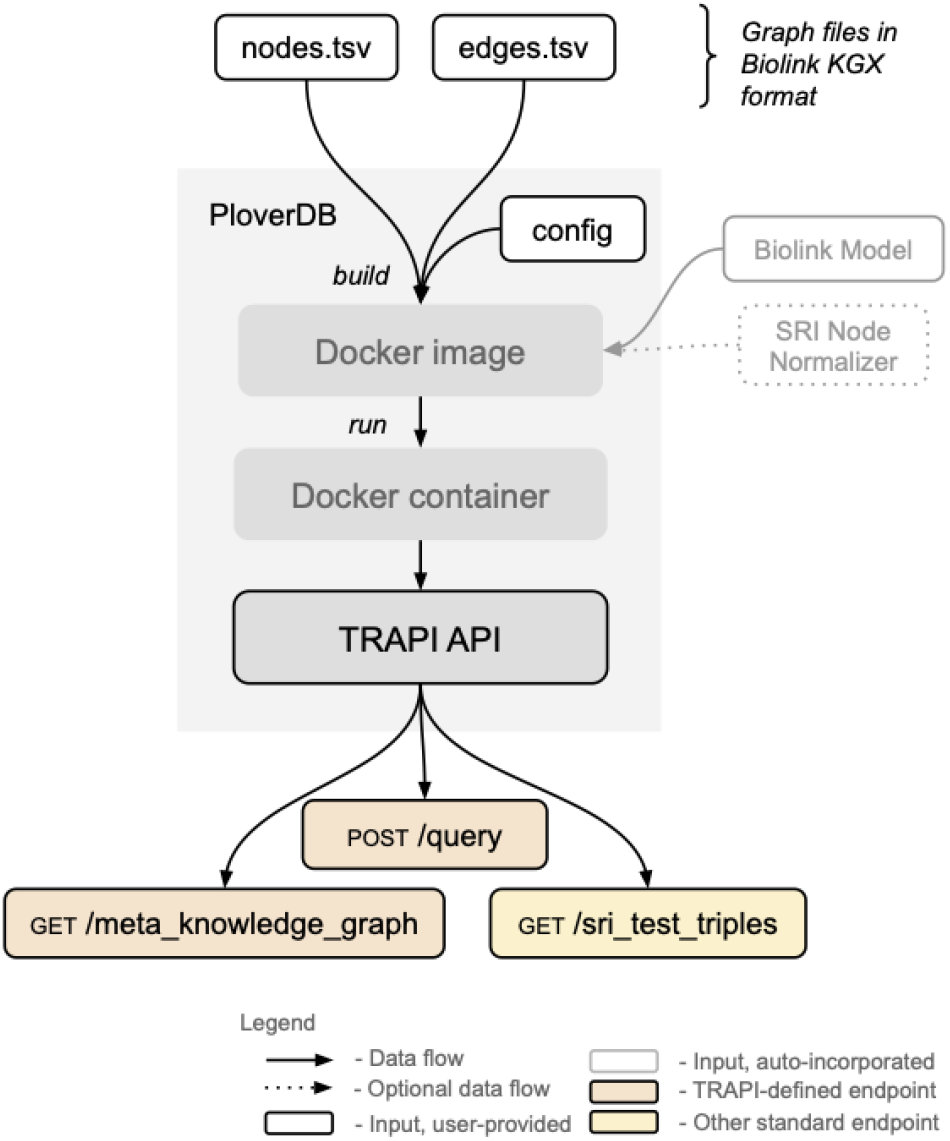
PloverDB’s high-level architecture, depicting how an input KGX-formatted knowledge graph is exposed as a TRAPI web API. Details are described in the main text.

### 2.1 Capabilities

PloverDB is designed to be a comprehensive platform for transforming a Biolink-compliant knowledge graph into a fully-functioning TRAPI-compliant API. Its capabilities in terms of supported queries (Sec. 2.1.1), built-in reasoning (Sec. 2.1.2), ease of deployment (Sec. 2.1.3), and performance (Sec. 2.1.4) are detailed below.

#### 2.1.1 Query structure

PloverDB is designed to answer one-hop TRAPI queries, which are essentially graph-based pattern matching queries consisting of two nodes connected by a single edge^2^. In such a “query graph”, nodes can be constrained by their concept (i.e., identifier) and semantic type (i.e., category) and edges can be constrained by their semantic type (i.e., predicate) and attributes (e.g., supporting publications, knowledge source, source type, etc.) Examples of such query graphs are provided in Appendix A.

#### 2.1.2 Built-in reasoning

Importantly, when PloverDB answers the queries defined in Section 2.1.1, it returns all knowledge subgraphs that fulfill the query graph not only *directly*, but also according to:

- the *hierarchies* of node and edge types according to Biolink,
- the *symmetry* of edge types according to Biolink,
- the *canonicity* of edge types according to Biolink, concept *equivalency*, and
- concept *subclass relationships* (via transitive chaining) present in the underlying graph or in a user-specified external source.

Of note, concept equivalency is determined either by 1) identifiers provided in an `equivalent_identifiers` (or similar) property on each node in the KG, or, in the absence of such a property, 2) the Translator Node Normalizer service. PloverDB fulfills most of these reasoning tasks at build time and encodes them into its indexes for constant-time lookup at query time.

#### 2.1.3 Simplified deployment

PloverDB is designed to streamline deployment so that data owners can easily and independently update their service upon KG content/representation changes. Thus, it:

1. is fully Dockerized and uses a standard Debian Linux-based base image from the public DockerHub registry (`tiangolo/uwsgi-nginx-flask:python3.11`),
2. automatically constructs and exposes a TRAPI meta knowledge graph and corresponding test triples,
3. allows multiple KGs to be served from the same PloverDB application, accessible at separate sub-endpoints (i.e., “multiplexing” of KGs),
4. provides a built-in (authenticated) remote deployment mechanism, enabling data owners to remotely redeploy their service with minimal downtime,
5. provides endpoints to aid remote debugging, that expose log and KG/code version information, and
6. can (optionally) be registered in SmartAPI [10] using the TRAPI registration template (github:Translator/ReasonerAPI/TranslatorReasonerAPI.yaml).

#### 2.1.4 Performance

PloverDB uses a fully in-memory architecture for efficient query answering. To quantify its performance, we conducted a controlled study comparing PloverDB to Plater, a previously-mentioned functionally similar platform that uses Neo4j for its back-end, when hosting the RTX-KG2 knowledge graph. The study included 82 queries for which the platforms returned successful responses: 79 real-world, randomly-selected queries and three hand-crafted queries, designed to address gaps in terms of answer size. We present highlights from this study below; full methodology details, results, and discussion for this study are provided in Appendix B.

In this controlled performance study, both platforms’ response times appeared asymptotically linear in terms of the number of edges in the query answer, but Plater had a three-times steeper slope than PloverDB, at 0.022 vs. 0.007 seconds per 100 answer edges. Figure 2.2 visualizes these results on a log-log scale; the magnitude of difference in query speed between the two platforms was greater for smaller queries, suggesting that Plater has higher overhead. The proportion of time Plater spent on Neo4j (as opposed to “wrapper” code around database calls) ranged from 80% to 99%, and showed a negative relationship with query answer size (Fig. B.7), suggesting Plater’s overhead arose from Neo4j.

PloverDB was 14-times faster than Plater on average across the 51 randomly-selected queries for which the two platforms returned identical answer sets (Table B.1). In a ramped load test to extreme conditions, PloverDB averaged 4.5-times higher throughput and 9-times faster response time than Plater (Sec. B.2.4).

PloverDB’s speed and low overhead observed in this study come with a trade-off; its in-memory design means that it is relatively memory-hungry, using 79 GiB of system memory at rest when hosting RTX-KG2.8.4c (7 million nodes and 27 million edges) vs. only 5 GiB for Plater. While PloverDB’s memory consumption depends heavily on the hosted graph’s content (e.g., the size and number of node/edge attributes) and configuration choices, our observations from this study and real-world deployments suggest that PloverDB tends to use roughly 3-5 GiB of memory per million edges.

## 3 Usage

Full technical details on how to use and deploy PloverDB are provided in GitHub at github:RTXteam/PloverDB. Assuming that Docker has been installed on the host computer and the PloverDB code repository has been cloned into the local file system, deploying PloverDB is simply a matter of:

1. editing `PloverDB/app/config.json` for the user’s KG, and
2. running `bash PloverDB/run.sh`, which builds and starts the Dockerized PloverDB service.

**Figure 2.2:**
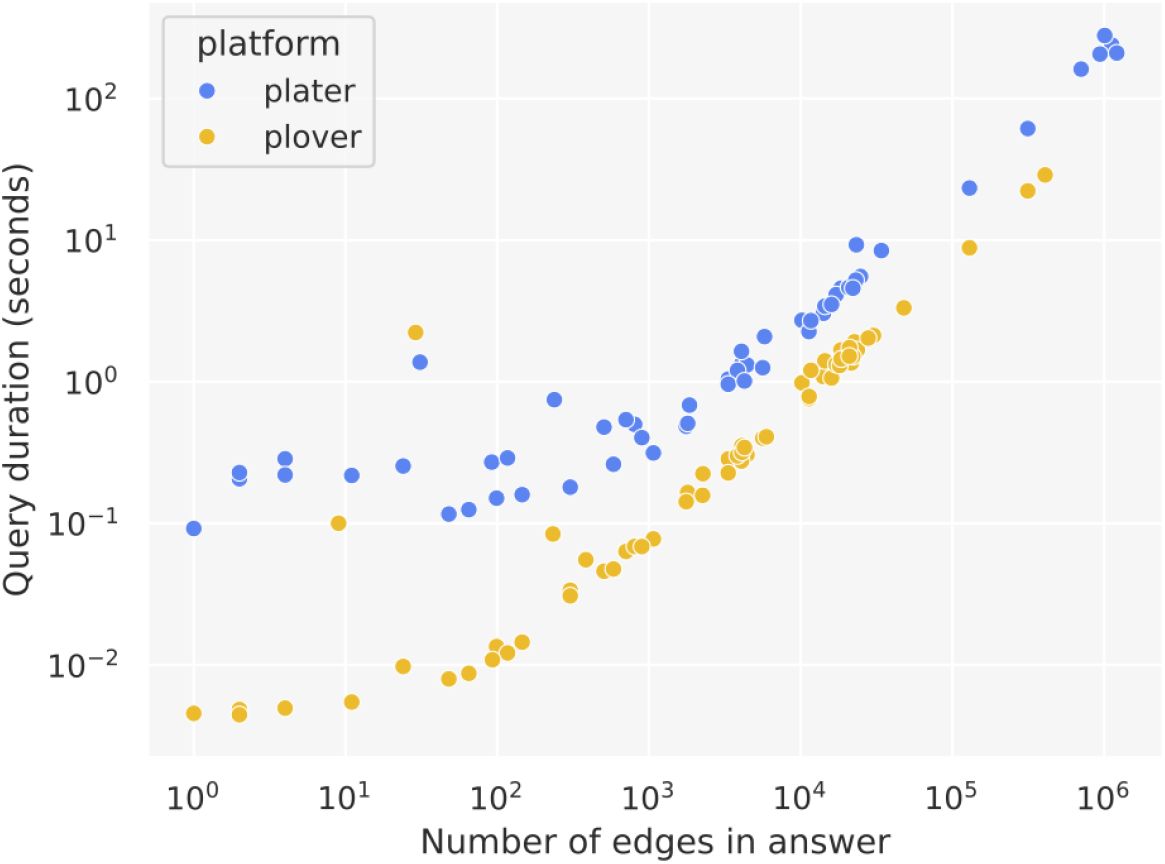
Query response time as a function of answer size for PloverDB vs. Plater when hosting the RTX-KG2 knowledge graph, based on a set of 82 largely randomly-selected real-world queries. Results are displayed on a log-log scale for better visibility; on a linear scale, both platforms form straight lines, but Plater has a three-times steeper slope than PloverDB (see Figure B.2). Details on the included queries are provided in Appendix Sections B.1.3 and B.2.1.

The PloverDB repository includes a template for the `config.json` file. Most significantly, users must update the `nodes_file` and `edges_file` slots with URLs from which PloverDB can download the graph’s nodes and edges files at image build time. These files must adhere to Biolink KGX (github:biolink/kgx) TSV or JSON Lines format. Users may also specify how node/edge properties should be represented as TRAPI Attributes using the `trapi_attribute_map` slot. Explanations for all `config.json` properties are available in the PloverDB GitHub repository, as well as steps to deploy PloverDB in a production setting, including TLS certificate installation.

The fact that the `config.json` file is tracked in the PloverDB software repository makes configuring continuous deployment straightforward; users can set up continuous deployment on a fork of the repository using a third-party tool, or they can use PloverDB’s built-in remote deployment mechanism, which allows users to deploy changes by submitting an authenticated `/rebuild` request to the PloverDB web service. Multiplexing KGs is a matter of adding an additional JSON config file for each KG to be served. PloverDB will automatically build and expose a TRAPI endpoint for each such KG, accessible at the `endpoint_name` specified in its configuration file.

At the time of writing, PloverDB is used to host and serve five different knowledge graphs for the Translator system: RTX-KG2 [9] (kg2cploverdb.transltr.io/kg2c), Microbiome KG [4] (multiomics.transltr.io/mbkp), Multiomics KG [7] (multiomics.transltr.io/mokp), Clinical Trials KG [7] (multiomics.transltr.io/ctkp), and Drug Approvals KG [7] (multiomics.transltr.io/dakp), ranging in size from 5,000 nodes and 45,000 edges to 7 million nodes and 27 million edges.

## 4 Conclusion

PloverDB is a software platform for efficiently hosting and serving Biolink-compliant knowledge graphs as standardized TRAPI-compliant web APIs. PloverDB emphasizes simplicity and automation, allowing knowledge graph owners to focus on their data rather than the technicalities of exposing it. Its comprehensiveness in terms of built-in reasoning and ease of deployment significantly reduce the barrier of entry for exposing knowledge graphs for use by others, thereby enhancing their utility for advancing biomedicine.

## Author contributions statement

A.K.G., S.A.R., and E.W.D. conceived the application. A.K.G. designed, implemented, and analyzed the software.

A.K.G. and S.A.R. wrote the manuscript.

## Acknowledgments

We thank David Koslicki, Gwênlyn Glusman, Mohsen Taheri, Kevin Vizhalil, Sundareswar Pullela, Skye Goetz, Chunyu Ma, Lindsey Kvarfordt, E. C. Wood, Evan Morris, and Max Wang for advice, inspiration, and/or feedback. We thank the NCATS Information Technology Resources Branch (ITRB) for incorporating PloverDB into their automated build and deployment system.

## A Examples of PloverDB queries and reasoning

To better illustrate the “one-hop” Translator Reasoner API (TRAPI) queries described in the main text that PloverDB is designed to answer, we provide some example query graphs below. They are represented in an abbreviated form, rather than full TRAPI JSON^3^.

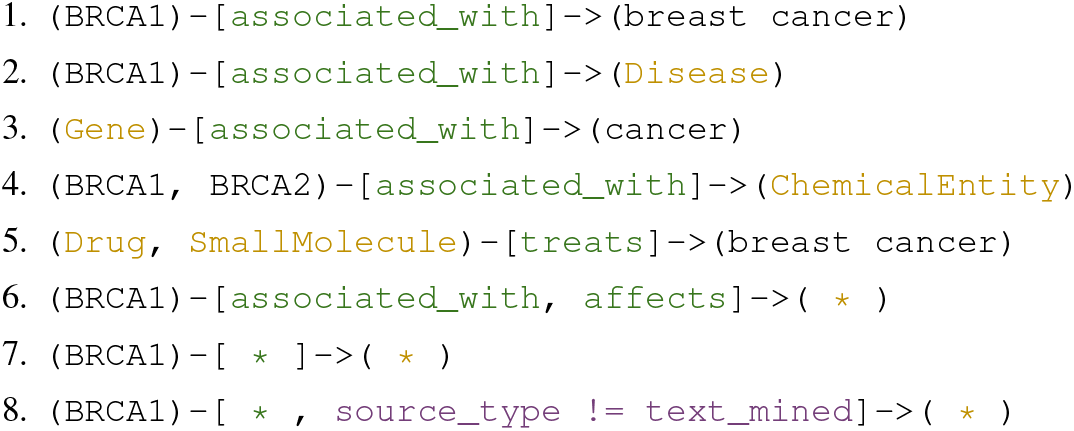

As explained in the main text, when answering such query graphs, PloverDB performs some knowledge reasoning tasks beyond direct pattern-matching. To better explain these, we have included a few examples below (based off of the above example queries).

- *Predicate symmetry/asymmetry*: When answering example Query 1, PloverDB ignores edge direction (i.e., it effectively also looks for patterns in the opposite direction), since the edge type `associated_with` is annotated as symmetric in the Biolink model. When answering Query 5, however, it will obey edge direction, since `treats` is an asymmetric predicate according to Biolink.
- *Concept subclasses*: When answering Query 3, PloverDB returns genes associated with the general concept of “cancer” as well as genes associated with any subclasses of cancer (e.g., breast cancer, pancreatic cancer, etc.).
- *Node category hierarchy*: When answering Query 4, PloverDB will return any chemical entities associated with *BRCA1* or *BRCA2*, as well as any small molecules, drugs, and other nodes with categories that are descendants of `ChemicalEntity` in the Biolink Model, that are associated with either of those genes.
- *Edge predicate hierarchy*: When answering Query 1, PloverDB will look not only for edges linking *BRCA1* and breast cancer that have the type `associated_with`, but also any descendant of that edge type in the Biolink Model, such as `correlated_with`, `gene_associated_with_condition`, etc.

While a full description of how TRAPI queries should be constructed is available in the TRAPI GitHub repository (github:NCATSTranslator/ReasonerAPI), we provide an example of a TRAPI query expressed in full JSON format below. This query submits our example Query 2 to the PloverDB-hosted RTX-KG2 API.

~~~
curl -X ‘POST’ \ ‘https://kg2cploverdb.transltr.io/query‘ \
    -H ‘accept: application/json’ \
    -H ‘Content-Type: application/json’ \
    -d ‘{
    “message”: {
       “query_graph”: {
        “nodes”: {
         “n00”: {
          “ids”: [
           “NCBIGene:672”
          ]
         },
          “n01”: {
           “categories”: [
            “biolink:Disease”
        ]
      }
     },
        “edges”: {
         “e00”: {
          “subject”: “n01”,
          “object”: “n00”,
          “predicates”: [
            “biolink:associated_with”
          ]
         }
        }
       }
      }
}’
~~~

## B Controlled performance study: PloverDB vs. Plater

This section details the controlled performance study behind the results presented in Section 2.1.4 of the main text. In this study, we compared PloverDB to Plater (github:TranslatorSRI/Plater), which uses Neo4j for its back-end database platform, and is the most functionally similar platform to PloverDB to our knowledge. We compared the performance of Plater vs. PloverDB when hosting the RTX-KG2 knowledge graph in terms of query speed (Sec. B.2.2), amortized query speed as a function of batch size (Sec. B.2.3), and throughput/load handling (Sec. B.2.4). We also compared system memory consumption, disk space consumption, and build time for the two platforms (Sec. B.2.5). We hypothesized that PloverDB would have faster query times and higher throughput due to its fully in-memory design, while Plater would use less system memory. We refer to PloverDB simply as “Plover” in this section, for (slightly) better readability.

Of note, we chose to compare the two software stacks (Plover and Plater) in their entirety, as opposed to only their back-end database platforms, in order to provide the most practical and actionable insight for knowledge graph owners choosing a hosting platform.

### B.1 Methods for controlled performance study

#### B.1.1 Platform and hosting details for Plater and Plover

The Plater and Plover platforms were hosted on separate `r5a.4xlarge` designated Amazon Elastic Cloud Compute (EC2) instances (128 GiB system memory, 16 virtual CPUs) cloned from the same base instance, running the Ubuntu Linux 18.04 operating system. Both hosting instances had 1 TiB of disk space in the root file system and were reserved strictly for our testing purposes. To best simulate real-world usage, test queries were sent via HTTP requests (without caching) from a third EC2 instance (also an `r5a.4xlarge`) in the same Amazon Web Services (AWS) Region as the hosting platforms (`us-west-2`), which we refer to as the “testing instance”. Both platforms spoke Translator Reasoner API (TRAPI) (github:NCATSTranslator/ReasonerAPI) version 1.4 and were configured with a 10-minute query timeout.

We used version 1.5 of Plater (which uses Neo4j version 4), with relatively minor modifications^4^. These included modifying Plater’s Neo4j Cypher queries to 1) perform multi-hop (instead of single-hop) concept subclass reasoning and 2) add a limit to the number of results^5^. We set the max depth for subclass chaining to 21 and the max number of results allowed to 1,000,000 for all tests, to align with Plover’s behavior. In regards to the first modification listed, it is worth mentioning that Plater v1.5 assumes that the knowledge graphs it ingests have materialized direct `subclass_of` edges between all pairs of nodes connected by a `subclass_of` *chain*; thus its Neo4j Cypher queries need only look for one-hop `subclass_of` edges to perform recursive subclass reasoning. We attempted this subclass prematerialization strategy with RTX-KG2.8.4c, but opted not to proceed with it because it did not appear to lead to faster query times on average in preliminary testing and it is unclear what proper provenance would consist of for the materialized `subclass_of` edges.

We used the Plover software as it existed in April 2024, captured at github:amykglen/PloverDB/releases/tag/v2.0. Of note, this version (and later versions) of Plover deliberately returns an error if a submitted query will produce an answer graph with more than 1,000,000 edges. In addition, neither platform performed entity resolution of input query node identifiers in this study (Plater does not provide such functionality, and Plover did not yet provide it at the time of the study).

#### B.1.2 Build processes used for Plater and Plover

The canonicalized version of RTX-KG2.8.4 was loaded into both platforms and used for all tests. Both platforms ingested a JSON Lines representation of the graph, which includes one file containing nodes and another containing edges. The edges file used was modified to remove two kinds of edges that Translator knowledge providers are currently not allowed to return per Translator architectural guidelines: 1) edges from SemMedDB with fewer than four supporting publications and 2) edges with invalid domain and range categories according to Biolink^6^. This means that the RTX-KG2.8.4c JSON Lines edges file that both platforms ingested contained about 27 million edges. Plater also requires that node categories are pre-expanded to their Biolink ancestors (up to `NamedThing`) in the knowledge graph it ingests, so an additional property was added to the nodes JSON Lines file to capture this. There were a total of 6.8 million nodes in the RTX-KG2.8.4c JSON Lines nodes file. Because our current RTX-KG2 API only performs subclass reasoning based on edges from a few trusted sources, we edited the RTX-KG2.8.4c JSON Lines file so that `subclass_of` edges from untrusted sources were assigned the higher-level predicate `related_to_at_ concept_level`, thus excluding them from concept subclass reasoning on both platforms.

To build Plater, we ran its back-end Neo4j database in a Docker container and the actual Plater application on the host machine. The script used to build Plater is available in GitHub at github:amykglen/plater-plover/run-plater-kg2.sh. We used ORION (github:RobokopU24/ORION) to load the RTX-KG2 JSON Lines file into Neo4j and create appropriate indexes.

To build Plover, we used its `run.sh` script (github:amykglen/PloverDB/blob/v2.0/run.sh) to generate a Docker image containing Plover and all of its necessary indexes (built from the RTX-KG2 JSON Lines files) and run a Docker container off of that image (which loads the indexes into memory).

#### B.1.3 Selection of test queries for controlled performance study

To generate a sample of real-world queries on which to compare the two platforms, we randomly selected a set of 90 queries out of those submitted to live (publicly-available web service endpoints registered via SmartAPI) RTX-KG2 knowledge provider services (which include development, staging, test, and production instances) during the prior three months: Dec. 6, 2023 to March 6, 2024. To cover a variety of RTX-KG2 use cases yet ensure queries to production instances (which are relatively less common) are still represented, we divided this sampling into three subsets, which we will refer to as indicated:

- “**Any**”: 30 queries randomly selected from the overall set of 11,234 unique^7^ RTX-KG2 queries submitted to any RTX-KG2 instance (development, staging, etc.) during the 3-month window
- “**ITRB prod**”: 30 queries randomly selected from the subset of 59 unique queries sent to the production RTX-KG2 instance hosted by the NCATS Information Technology Resources Branch (ITRB) during the 3-month window
- “**Long**”: 30 queries randomly selected from the subset of 351 queries with an elapsed time over 30 seconds during the 3-month window

For the first two of these samplings, we excluded queries with an elapsed time^8^ under two seconds to bias the sample towards queries that actually produce answers (answer size is not recorded for past RTX-KG2 queries), since we are most interested in analyzing speed for queries that RTX-KG2 can actually answer. In addition, we included three hand-crafted queries designed to produce very large answer sets in comparison to typical queries of RTX-KG2 (i.e., *>*300k edges) to supplement the analysis of query speed as a function of answer size in Section B.2.2; these hand-crafted queries were not included in any other analyses. The final 93 queries are available on GitHub at github:amykglen/plater-plover/tree/main/test. Testing was performed in April 2024.

We forced `is_set=False` on all query nodes in all test queries, as this represents the most realistic real-world scenario for queries of RTX-KG2^9^.

#### B.1.4 Testing and data collection methods for controlled performance study

For the main analysis of query speed (Sec. B.2.2), we ran three repetitions of the full test suite of 93 queries against each platform. Queries were run in random order on each run of the test suite^10^. Between those three runs, the hosting instances were both rebooted and the platforms were rebuilt to clear any cached data (in the operating system or elsewhere). The testing instance was also rebooted between the three runs. Query duration was measured as elapsed wall clock time from the beginning of the query until the response *began* to be streamed back to the testing instance (i.e., when the response headers were returned).

For the amortized analysis of query speed as a function of batch size (Sec. B.2.3), we selected the longest-running query with more than 1,000 pinned node IDs that produced near-identical answer sets from the two platforms (out of the total 90 randomly-selected queries described above). This query was sent sequentially to each platform with batch sizes of 1, 10, 100, and 1000. Here a “batch size” refers to the number of pinned node identifiers; a batch size of one means that we sent 1,894 copies of the query, each with only *one* of the pinned node IDs; a batch size of 1,000 means that we sent two copies of the query, one with 1,000 of the pinned node IDs and the other with the remaining 894 node IDs.

Load testing (Sec. B.2.4) was performed using the Python package Locust (docs.locust.io) (v2.26.0). For both platforms, an 18-minute ramped load test was conducted in which the number of simulated “users” was increased from 0 to 2,000 at a spawn rate of 2 users per second. Each user submitted one query every 5 to 20 seconds (as opposed to at a fixed interval) in order to better simulate real-world behavior of reasoning agents and other users of RTX-KG2. This means that, on average, simulated users submitted a query every 12.5 seconds; thus, 2,000 concurrent simulated users was equivalent to an average of about 160 queries per second^11^. Simulated users always randomly selected which query to send out of the set of “ITRB prod” test queries, excluding those for which 1) either of the platforms returned a non-`200` HTTP status code or 2) the two platforms returned differing answer sets in the main analysis of query speed (Sec. B.2.2). To analyze the results, throughput rates, response times (min, max, and average), and number of simulated users at each second of the load test were extracted from the `stats_history.csv` file that Locust outputs. One run of the load test was performed per platform. The testing instance was rebooted between load tests.

The system memory usage amounts reported in Section B.2.5 were obtained using the `psutil` (psutil.readthedocs.io) Python library’s `virtual_memory` method; a background process repeatedly called this method (every 10 seconds during platform builds and every 1 second during batch querying tests) and recorded the percentage of memory used according to the returned `percent` value. Percentages were then converted to GiB values by calculating (*percent/*100) ∗ *total*, where `total` is the total physical memory according to `psutil.virtual_memory` (124.7 GiB for both hosting instances). Memory usage during builds was averaged across three build repetitions for each platform.

For Plater, the disk space usage amount reported in Section B.2.5 necessary to run the platform were calculated by adding the sizes of the `Plater` and `neo4j` directories^12^ on the hosting instance. The additional disk space required during the Plater build was the size of the `ORION_parent_dir` directory on the hosting instance. For Plover, the disk space necessary to run the platform consisted of the size of the outer `app` directory in the Plover Docker image produced during the build, which directly includes all of Plover’s indexes and necessary files (not mounted as volumes). The additional disk space required during the Plover build corresponds to the size of the unzipped RTX-KG2 JSON Lines nodes and edges files.

The build times reported in Section B.2.5 were calculated as wall clock time from the initiation of a fresh build process (without caching) to the point when the platform was fully up and running and ready to receive queries. This can be thought of as the time to go from the underlying graph data in JSON Lines files to a live TRAPI API serving that graph. Build times were averaged across three repetitions.

Analyses were done in Python using Pandas (v2.0.3) and SciPy (v1.11.4) and in Google Sheets. Statistical tests were performed using SciPy’s `stats` module.

### B.2 Results for controlled performance study

#### B.2.1 Final included queries

Interestingly, despite being loaded with exactly the same knowledge graph and adhering to the same built-in reasoning tasks, Plater and Plover only returned identical answer sets^13^ for 69% of the total 93 test queries; 17% of the test queries only produced answers from *one* of the platforms and the remaining 14% of queries produced partially-overlapping answer sets. We considered answer sets to “match” for our analyses if the number of edges differed by *<*1% between the two platforms; 69 of the total 93 queries met this criteria.

Of the 69 matching queries, only 52 returned 200 status codes from both platforms. Potential reasons for these differences are discussed in Section B.3. To ensure a fair comparison between the two platforms, we only included these 52 successful, matching queries in our analyses, with the exception of the exploration into query speed as a function of answer size in Section B.2.2. For that analysis, we still included only successful queries, but kept the queries that produced differing answer sets since each platform’s durations are plotted as a function of *their own* answer size in this analysis. Note that hand-crafted queries were also only included in the analysis of query duration as a function of answer size, as described in Section B.1.3. In the end, a total of 82 queries were included in the analysis of query speed as a function of answer size and 51 queries were included in all other analyses of query speed.

#### B.2.2 Query speed results from controlled performance study

Results for the direct comparison of the two platforms on a per-query basis are visualized in Figure B.1, which shows that Plover outperformed Plater in terms of query duration on all 51 queries. On average across all queries, Plover was 14-times faster than Plater. A paired samples Wilcoxon signed-rank test showed a statistically significant difference in query durations between the two platforms (*W* = 169, *p-value <* 0.001). This result as well as the results for each of the three subsets of test queries are shown in Table B.1 below. Plover was significantly faster than Plater for all three subsets of test queries.

**Table B.1:** Summary of the average speedup provided by Plover over Plater for the various test subsets, including only queries that produced identical answer sets from the two platforms. The *n* column represents the number of queries in each subset. Average speedup was obtained by calculating the ratio of Plater’s mean duration to Plover’s mean duration for each query in the specified subset, and then averaging those ratios. Paired samples Wilcoxon signed-rank tests were performed between Plater’s and Plover’s sets of *n* (mean) query durations to obtain the *W* statistic and p-value for each test subset.

| Test subset | n | Average speedup | W | p-value |
| --- | --- | --- | --- | --- |
| Overall | 51 | 14-times | 169 | <0.001 |
| Any | 15 | 26-times | 12 | 0.004 |
| ITRB prod | 19 | 11-times | 22 | 0.002 |
| Long | 17 | 5-times | 20 | 0.006 |

<sup>12</sup>The `neo4j` directory on the hosting instance is mounted as a volume for the Neo4j docker container.
<sup>13</sup>Answer sets were considered identical if they returned the same *edges*; the number of TRAPI *results* differed somewhat, as the two platforms have slightly different (but both valid) ways of constructing results.

**Figure B.1:**
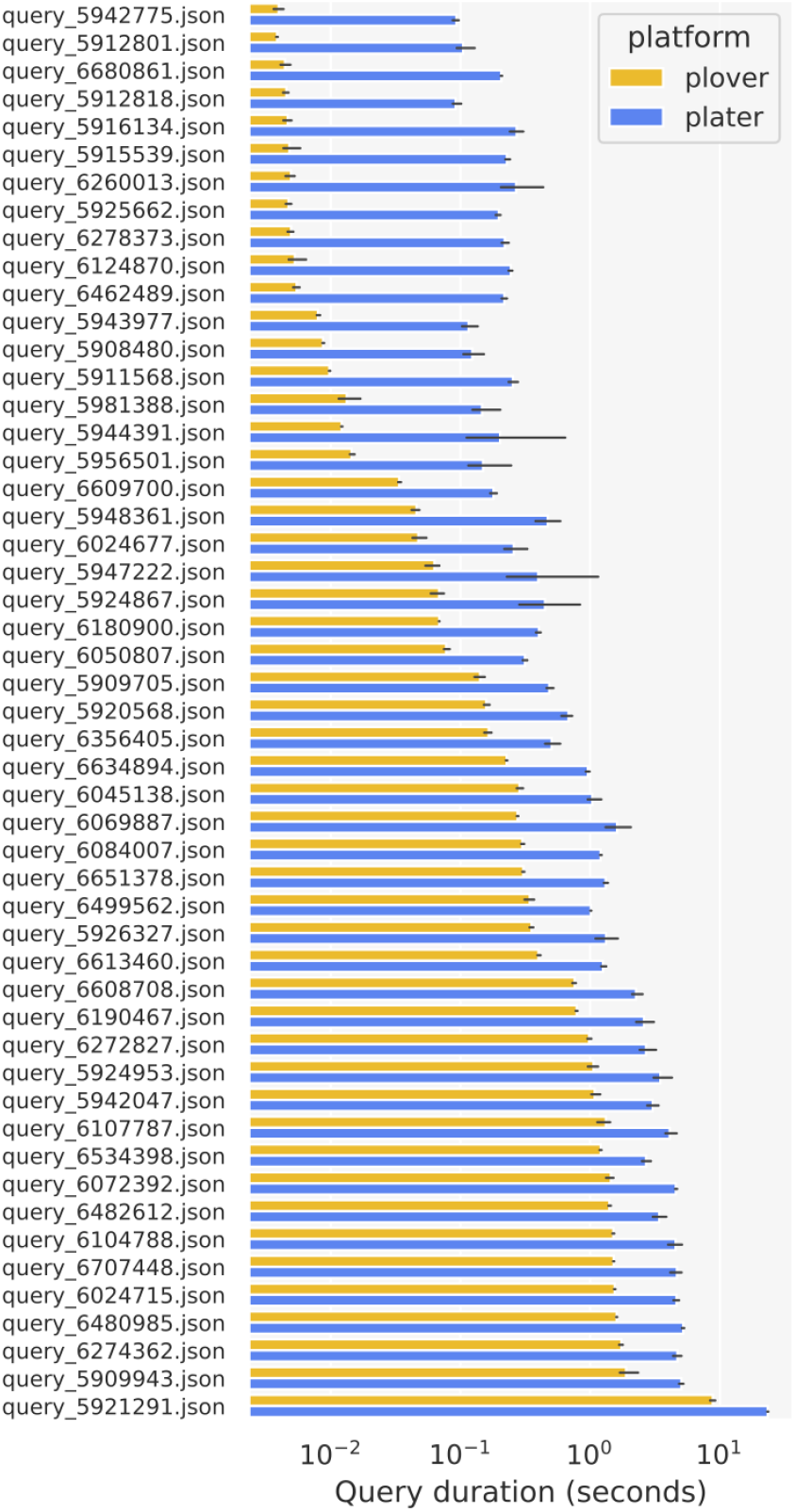
Mean query durations for Plater vs. Plover on a real-world sample of 51 queries. Whiskers (in black) indicate 95% confidence intervals. Note the log scale on the x-axis.

As can be seen in Figure B.1, the magnitude of difference between the two platforms appears to generally decrease with increasing query duration. This can be observed more clearly in Figure B.2, which visualizes query speed as a function of answer size. The asymptotic limit of the relationship between query duration and answer size in the top plot in Figure B.2 is **linear** for both platforms, with Plater’s slope three-times steeper than Plover’s, at 0.022 vs. 0.007 seconds per 100 answer edges. Plover appears to have lower overhead than Plater, with the magnitude of difference between the two platforms going from over 10-times for the shortest queries to roughly three-times for queries producing 100,000 or more answer edges.

Interestingly, Plater was found to have higher variability in query duration across the three test runs than Plover, with an average coefficient of variation^14^ of 0.16 vs. 0.06. This difference was statistically significant according to an unpaired samples Wilcoxon rank sum test (*U* = 3.4, *p-value <* 0.001). As can be seen in Figure B.3, there does not appear to be an obvious relationship between the two platforms in terms of their variability on each query.

#### B.2.3 Amortized query speed results from controlled performance study

The query selected for the amortized analysis (according to the criteria specified in Section B.1.4) was `query_5909943.json`, whose answer set sizes from the two platforms differed by *<*0.05%^15^. This query asks for drugs that are “part of” any of a number of specified concepts – i.e., `(1,894 node IDs)-[has_part]->(Drug)` – and was originally submitted by the ARAGORN (github:ranking-agent/aragorn) reasoning agent to the ITRB CI RTX-KG2 instance.

Amortized speed as a function of batch size for this query is shown in Figure B.4. Plater demonstrates a much greater decrease in amortized query speed as batch size goes up, which aligns with the observation that Plater appears to have more overhead than Plover in Section B.2.2. In addition, these data suggest that Plover reaches a lower asymptote than Plater as batch size gets larger, indicating that Plover is more efficient on a per-unit (i.e., per-pinned node ID) basis.

#### B.2.4 Load testing results from controlled performance study

Load testing was performed as described in Section B.1.4, essentially consisting of a ramp test in which the number of simulated concurrent users was increased at a rate of 2 per second to an extreme of 2,000 concurrent users. Each user submitted a query every 5 to 20 seconds, randomly selecting the query from the 19 “ITRB prod” test queries for which the two platforms both returned 1) 200 status codes and 2) identical answer sets during the main analysis of query speed (Sec. B.2.2). Importantly, the fact that we only included queries for which neither platform returned errors under normal conditions means we can assume that failed requests during the load test are a result of high load.

Throughput (successful requests per second) as a function of the number of simulated users throughout the ramp test is visualized in Figure B.5, which shows that Plover outperformed Plater. The points at which Plater and Plover began to regularly return failed requests are marked with blue and yellow arrows/vertical dashed lines, respectively; this threshold was at about 400 concurrent users for Plater and 780 concurrent users for Plover. This represents a load of approximately 32 vs. 62 submitted queries per second, respectively, given that each simulated user submitted a query at an average of every 12.5 seconds (as described in Section B.1.4). Interestingly, Plover’s throughput shows strong periodicity beyond the point at which it began to regularly return failed requests; potential reasons for this are discussed in Section B.3.

Plover also exhibited faster average response time throughout the load test, as can be observed in Figure B.6. The points at which Plater and Plover began to regularly return failed requests are again marked by blue and yellow arrows, respectively, in this figure.

Summarized results from the load test are reported in Table B.2. On average over the course of the ramped test, Plover had a 4.5-times higher throughput and 9-times faster response time than Plater. The percentage of requests per second that were successful out of total requests processed per second (i.e., failed + successful) averaged 58% for Plater vs. 71% for Plover over the course of the test to extreme conditions.

#### B.2.5 Space consumption and build time results from controlled performance study

Unsurprisingly given its in-memory design, the Plover platform used much more system memory than the Plater platform both during builds (max of 113 GiB vs. 6 GiB) and at rest^16^ (79 GiB vs. 5 GiB). Memory consumption beyond that at rest would be expected to vary depending on query load, but exploration of that relationship is beyond the scope of this study. However, we recorded memory usage levels during the batch testing performed for the amortized analysis (Sec. B.2.3) – which consisted of sequentially sending queries to the two platforms – and found that memory usage increased over the “at rest” level by a maximum of 0.4 GiB for Plover and 2 GiB for Plater.

**Figure B.2:**
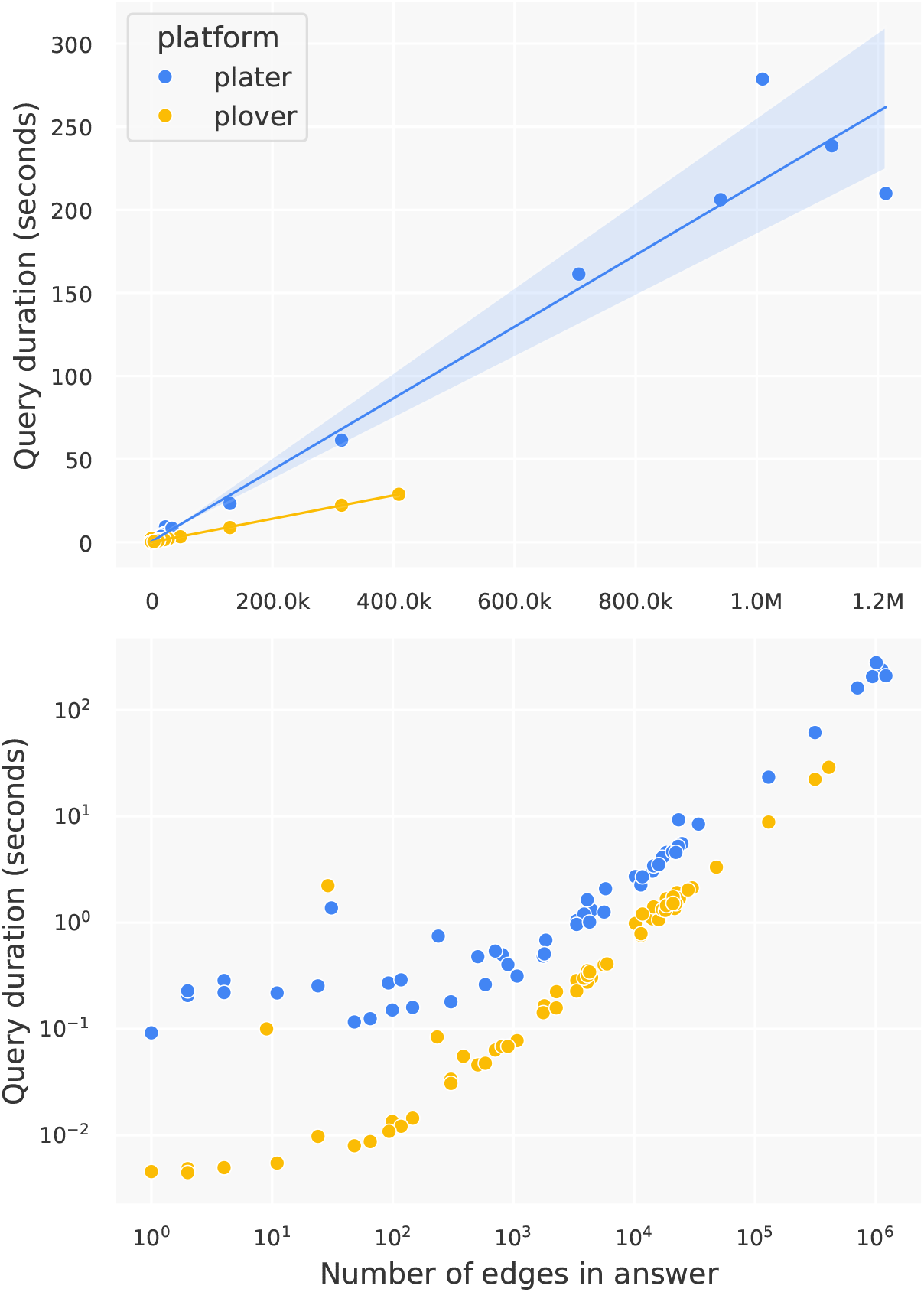
Query durations for Plater vs. Plover as a function of answer size on a linear scale (top) and log-log scale (bottom), for the 82 included queries for which the platforms returned successful responses. Both the x-axis and y-axis reflect the mean values across the three test runs. **Top plot**: The top plot shows least-squares linear regression trendlines, with shaded 95% confidence intervals (*R*^2^=0.971 and 0.996 for Plater and Plover, respectively). Both platforms appear to have asymptotically linear query durations, but Plover’s slope is about one-third that of Plater’s, at 0.007 vs. 0.022 seconds per 100 answer edges. Plater’s five largest queries – obvious on this linear scale – are queries that produced more than a million answer edges when using Plover, that Plover therefore deliberately returned an error for (this is its default behavior to avoid overwhelming clients). **Bottom plot**: The log-log scale illuminates the larger difference between the two platforms for smaller answer sizes, suggesting Plater has higher overhead than Plover.

**Table B.2:**
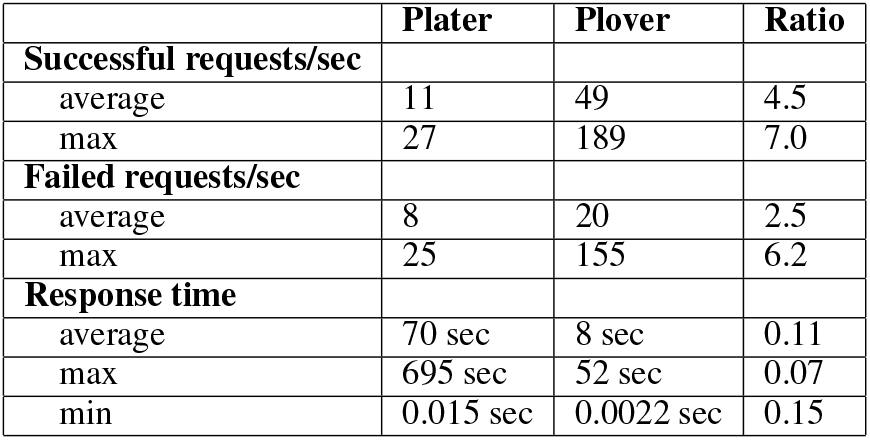
Summarized results for Plater vs. Plover during the ramped load test to extreme conditions. Statistics are the average/max/min of the measures that Locust reported at every second of the load test. Both platforms reached 2,000 users at the 1,001st second of the load test; the statistics below are based on the first 1,011 seconds of the test for both platforms. Each measure excludes the initial seconds of the load test prior to which data actually began to be reported for that measure, with the exception of average response time, which is the average response time Locust reported at the final timestep of the load test (the 1,011th second). The “Ratio” column reports the ratio of Plover’s result to Plater’s result. Abbreviations: “sec” refers to seconds.

**Figure B.3:**
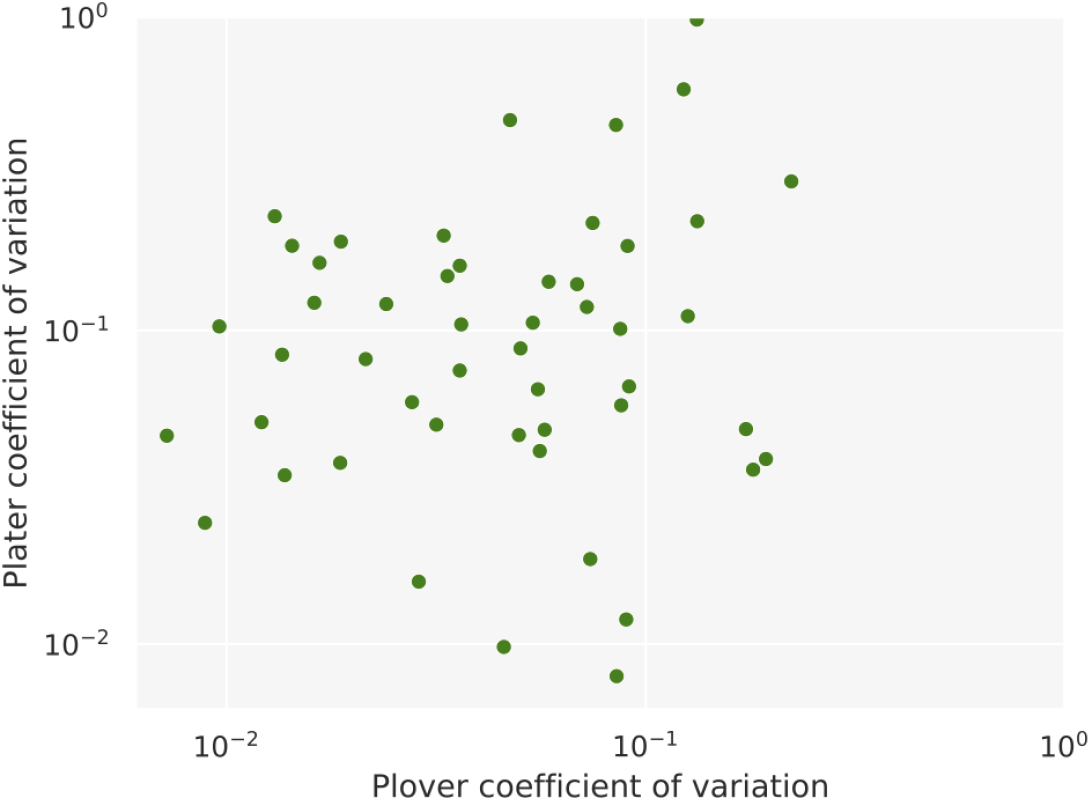
Each query’s coefficient of variation across the three test runs for Plater vs. Plover. Each dot in the scatterplot corresponds to one of the 51 included test queries. The coefficients of variation for the two platforms do not appear to be particularly correlated.

**Figure B.4:**
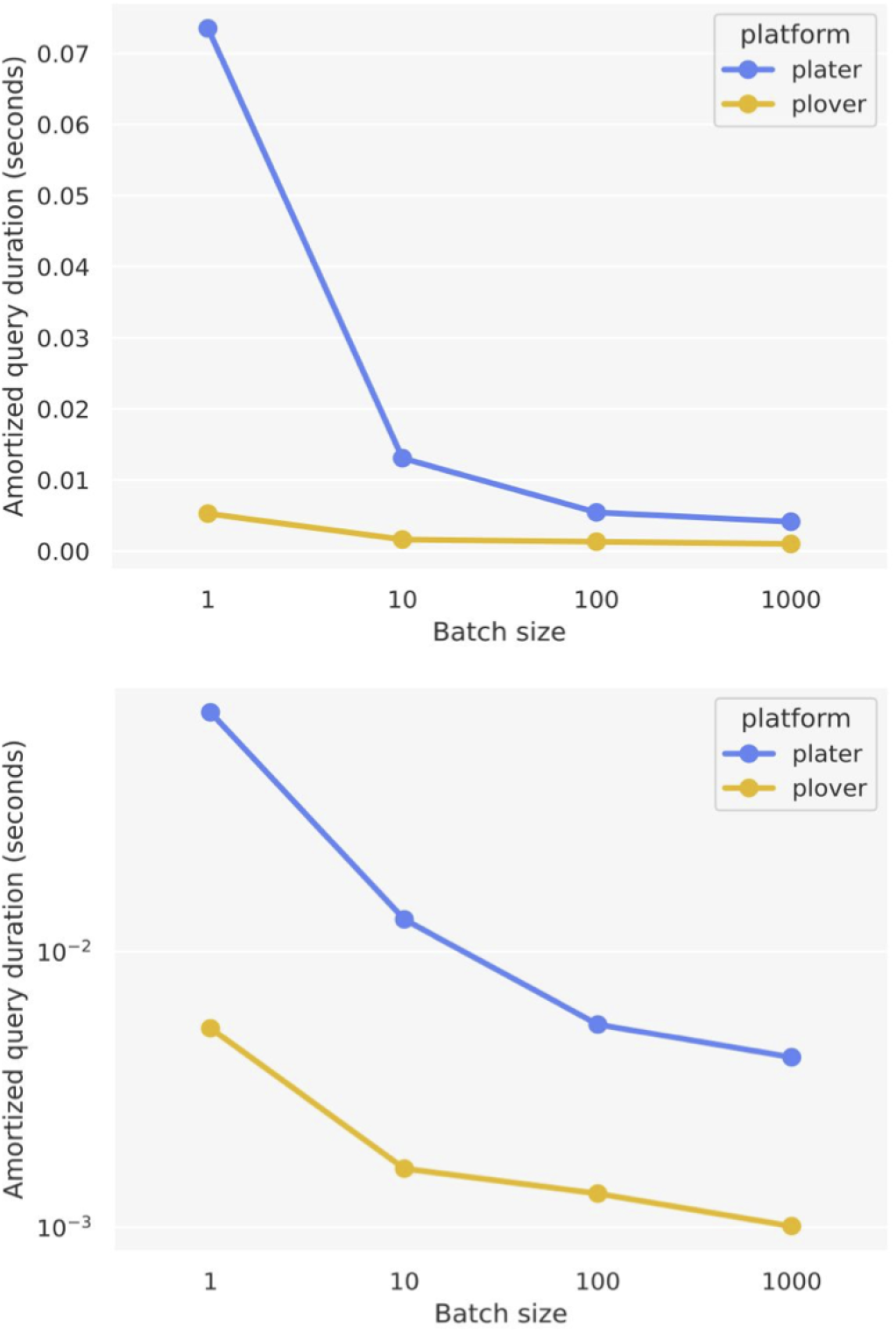
Amortized query speed as a function of batch size for Plater vs. Plover. The two charts differ only in the scale of their y-axes, with a linear scale in the top chart and a log scale in the bottom chart. Data is from a single test query (`query_5909943.json`), submitted using varying batch sizes (i.e. number of pinned node IDs). Amortized query duration is the wall-clock time taken to process a single pinned node ID – i.e., *(sum of query durations for a given batch size)/*1,894, where 1,894 is the total number of pinned node IDs in the test query.

**Figure B.5:**
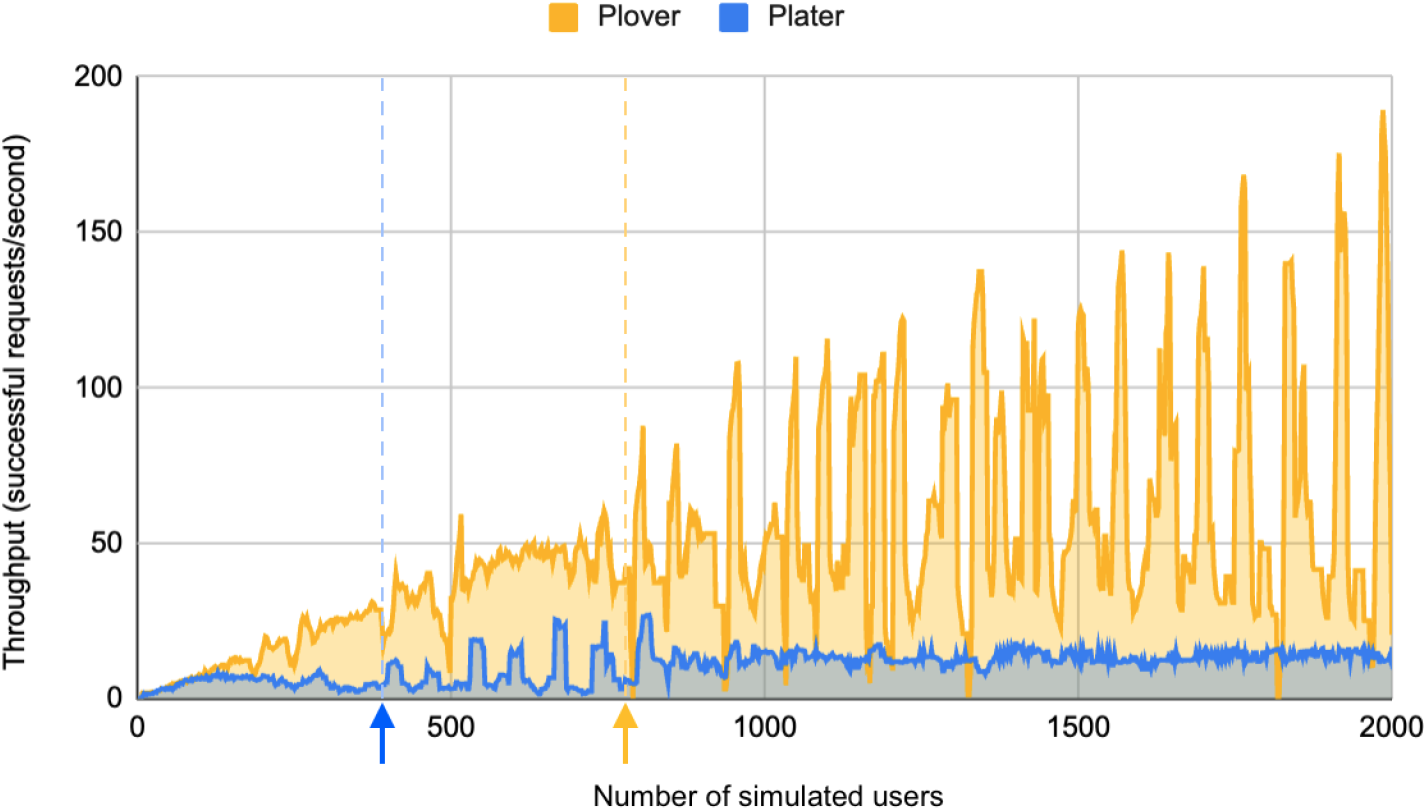
Throughput as a function of concurrent load for Plater vs. Plover. Area line charts are overlaid for the two platforms. The two blue and yellow arrows and corresponding dashed vertical lines mark the points at which Plater and Plover respectively began to regularly return failed requests. Note that since users were spawned at a rate of 2 per second, the x-axis can also be viewed as timesteps of the load test.

**Figure B.6:**
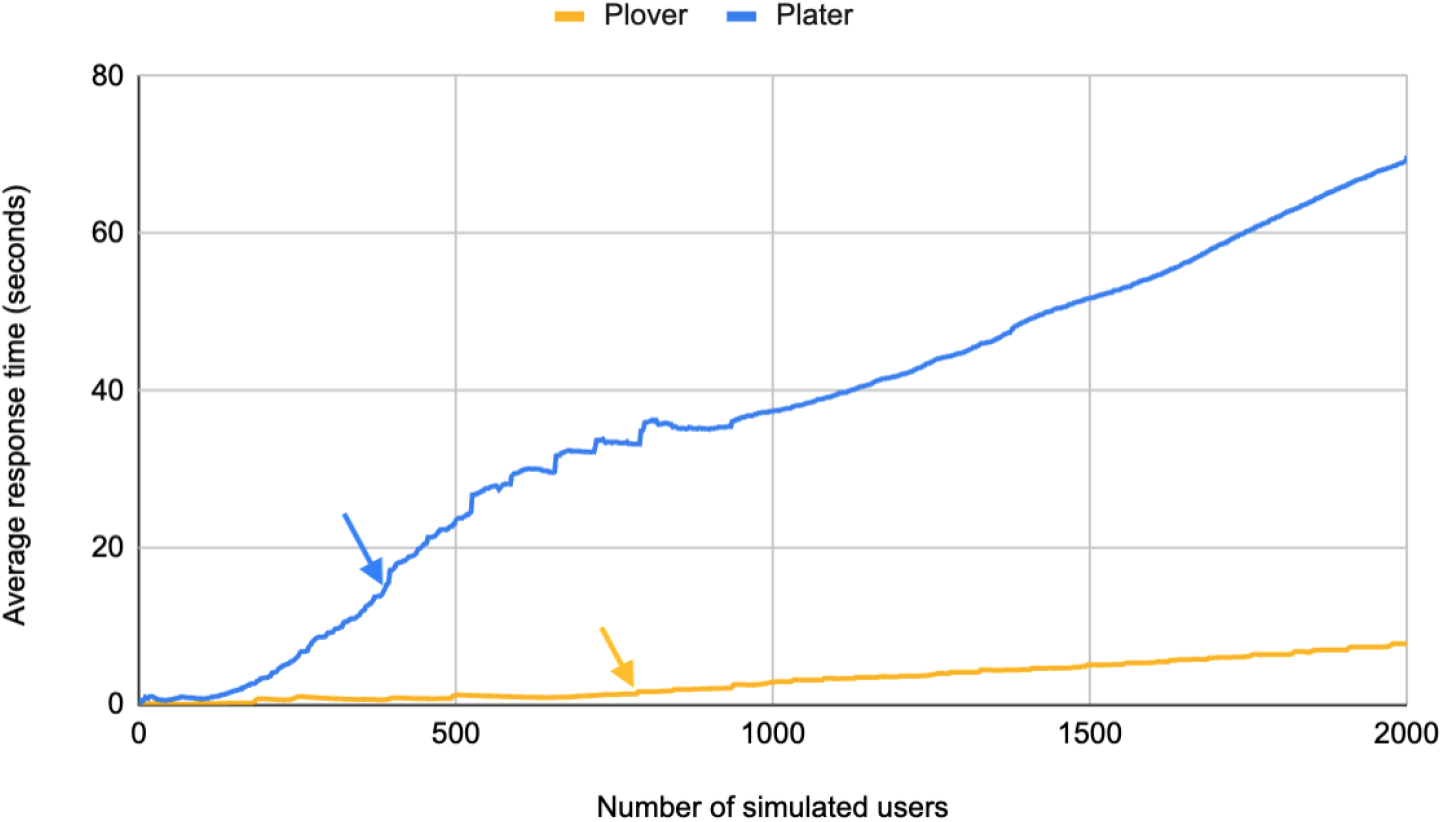
Average response time as a function of concurrent load for Plater vs. Plover. Average response time represents the *total* average response time up to that point in the load test (meaning, the average response time at the last timestep of the load test represents the overall average response time during the whole load test). The blue and yellow arrows indicate the points at which Plater and Plover respectively began to regularly return failed requests in the ramped test. Note that since users were spawned at a rate of 2 per second, the x-axis can also be viewed as timesteps of the load test.

The amout of disk space used by the two platforms was not drastically different, at about 23 GB for Plater vs. 17 GB for Plover. The Plater build process required an additional ≈ 35 GB of disk space for ORION, though those files do not need to be present to actually *run* Plater, as Plater relies only on the Neo4j dump that ORION creates. Plover similarly used more disk space during its build process, specifically about 18 GB to store the RTX-KG2 JSON lines files, which are then deleted at the end of the Docker image build. Thus when counting the overall disk space used while building *and* running the platforms, Plater’s total was 58 GB and Plover’s total was 35 GB.

As for build times, Plater builds were about 11-times faster than Plover builds, taking 5 minutes vs. 57 minutes to import the RTX-KG2 JSON Lines files and stand up a live and ready TRAPI API service. Of note, the Plater build requires that categories have been pre-expanded to their ancestor categories in the RTX-KG2 JSON Lines files, while Plover does not (it does this expansion during its build); however, we would not expect this to account for much of the difference in build time between the two platforms. Potential reasons for this difference in build times are discussed in Section B.3.

### B.3 Discussion of controlled performance study

Our results confirm our hypothesis that Plover would have faster query times and higher throughput while Plater would use less system memory. Supporting our theory that the performance difference between the platforms is primarily due to their back-end data stores (as opposed to “wrapper” code surrounding database calls), we found that Plater spent the majority of query time on its Neo4j calls (88% on average, ranging from 80% to 99%). Interestingly, as can be seen in Figure B.7, the percentage of time Plater spent on Neo4j appears to have somewhat of a negative relationship with query answer size, with queries producing *<*500 edges spending 94% of their time on Neo4j vs. queries producing *>*1,000 edges spending 84% of their time on Neo4j, on average. Given that the magnitude of difference in query speed between Plater and Plover *decreases* with increasing answer size, we believe this further suggests that the majority of the difference in speed between the two platforms may arise from Plater’s back-end database. Notably, while the version of Plater we used for these experiments used Neo4j 4, Plater was upgraded to Neo4j 5 after our study, which may result in improved processing times.

**Figure B.7:**
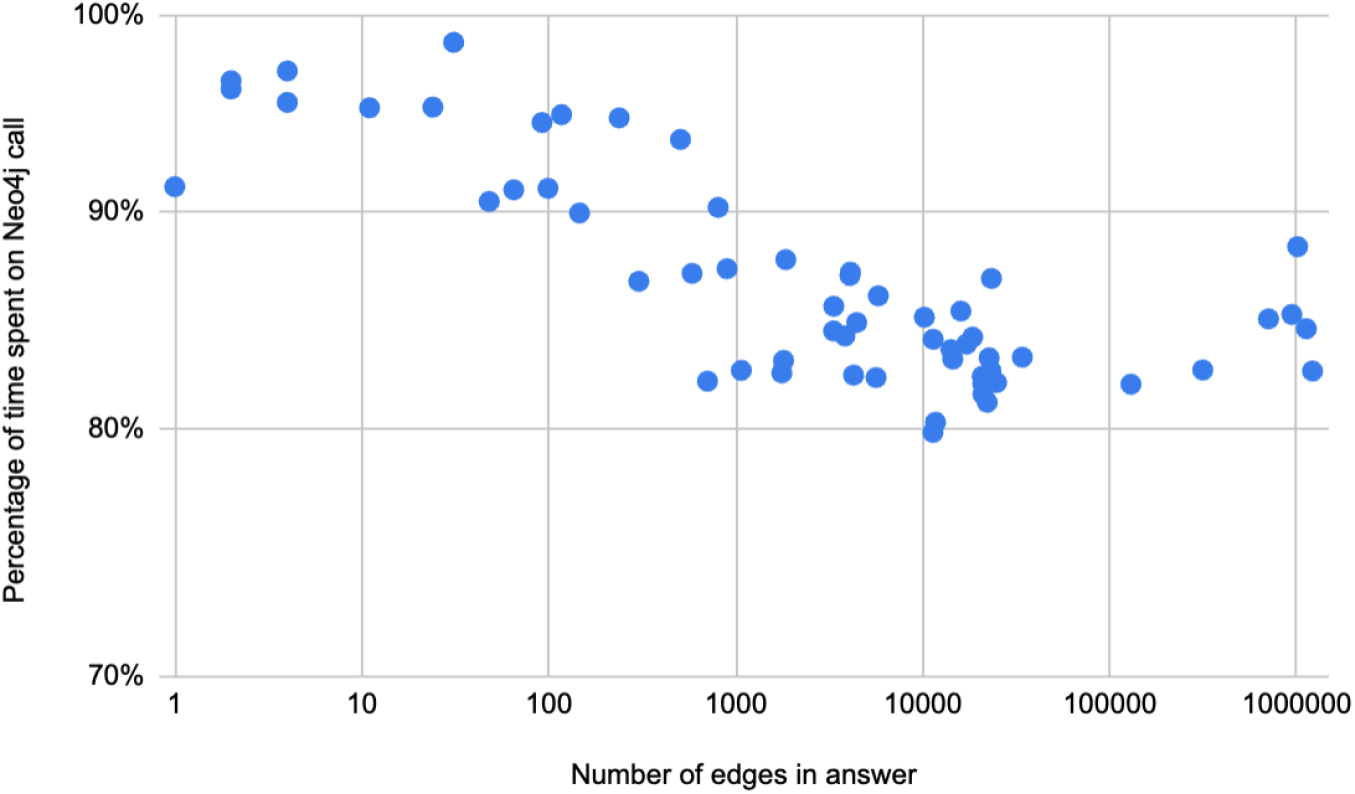
Percentage of time Plater spent on its Neo4j call out of the total duration of each query. Note that the y-axis begins at 70% in order to better visualize the change in percentage as answer size increases.

Further aspects of the results, potential limitations, and areas for future work are discussed in more detail in the below sections, organized thematically.

### Answer set differences

We can think of a couple potential reasons as to why the Plater and Plover platforms did not return identical answer sets for 31% of the original 93 selected test queries. Most significantly, while both platforms used a max depth of 21 for `subclass_of` chaining, Plover excludes nodes with more than 5,000 descendants from its subclass index in an attempt to avoid unhelpful/”overly general” nodes (such as the concept “biological products”) that stand to bloat answers. However, we do not believe this affects our findings since we excluded queries that produced differing answer sets from our analyses (with the exception of the analysis of query speed as a function of answer size, in which their inclusion is appropriate). Second, a number of the non-matching answer sets were for queries where Plater returned an error while Plover did not. This could be due to a bug in Plater, but investigating this was beyond the scope of this project.

### Load testing

The periodicity in Plover’s throughput during the ramped load test is curious, and we suspect it may be a result of the way in which Locust simulates concurrent users. During Plover’s load test, Locust output a warning shortly after reaching 2,000 users that indicated it was struggling to simulate such a high load using only one core. We attempted another run of the Plover load test that distributed the simulation across multiple cores on the testing instance (by specifying `-processes -1` in the Locust run command), but the `stats_history.csv` file that Locust saves test data to appeared to be subject to write conflicts between the parallel processes, leaving it unusable. Further load testing with improved user simulation methods would be worthwhile. However, despite this volatility in Plover’s throughput data, the present results still indicate that Plover is more scalable to a higher concurrent load than Plater, as evidenced by its higher threshold at which it began to regularly return failed requests and its higher average throughput vs. Plater.

In addition, our load testing results are likely overly optimistic as to Plater’s throughput and response times given that Plater uses caching and exhibits much faster query times for repeated runs of the same query, and our load testing essentially consisted of repeatedly sending the same 19 queries. We did not disable Plater’s caching in our testing because 1) it appears to be interwoven into Neo4j and the operating system in a way that is difficult to extricate and 2) the caching does not appear to be strictly on a per-query basis, meaning that caching may help improve query times even over runs of *different* queries, which represents an inherent speedup mechanism that is an advantage even in real-world situations where queries are *not* repeated, and thus it would be unfair to disable it. Caching should not have unfairly affected our other analyses of query speed outside of load testing since the platforms were fully rebuilt and the hosting instances rebooted between runs of the test suite, and the test suite did not contain duplicate queries. Plover did not exhibit noticeably faster processing of repeat queries in our experience, though we still rebuilt/rebooted it between test runs in the same way that we did for Plater.

Further load testing with inclusion of test queries beyond only the 19 included ITRB Prod queries to reduce the repetitiveness of queries may be worthwhile. In addition, the ITRB Prod queries tended to be quite small/short-running, but large queries (i.e., those producing *>*500k answer edges), while relatively infrequent, do happen in practice for our live RTX-KG2 APIs and it is important that such queries do not “take down” the system. Future load testing could incorporate such long queries as well.

### Variability in query times

Plater’s higher variability in query times across runs of the test suite in the main analysis of query speed in Section B.2.2 (despite clearing caches between runs) could be related to the fact that we randomized the order in which queries were run. Neo4j’s complex caching mentioned in the above discussion of load testing could mean that, even though there were no duplicate queries in each test run, similarities between queries such as overlap in their answer sets might result in certain kinds of caching that help Neo4j access needed data more quickly. Indeed, when breaking down our variability results further, we found that Plater’s Neo4j calls appear to account for most of its variability; on average across all queries^17^, Plater’s coefficient of variation strictly for time spent on Neo4j was 0.16 vs. 0.07 for time spent *not* on Neo4j, with the latter quite close to the variability observed with Plover (0.06). An in depth exploration into Neo4j’s caching methods is beyond the scope of this project, but we mention it here as a potential explanatory factor for this variability.

### Space consumption and build time

The difference in Plater and Plover’s build times is likely largely due to the fact that Plover creates 79 GiB-worth of in-memory indexes during its build process which it then both 1) saves to disk near the end of the Docker image build step and 2) loads back into memory during the Docker container run step. This represents a classic tradeoff in computer science where time spent on precomputation (i.e., building extensive indexes) allows for time savings down the road (i.e., faster query times). Given that RTX-KG2 is rebuilt no more than once every two weeks in practice, a build time of 1 hour is relatively trivial. Thus, while Plater’s quick 5-minute build time is convenient for development work, we do not see this as a significant advantage in terms of actual deployment of the platforms. Plater’s lower system memory usage, however, does allow for RTX-KG2 to be hosted much more cheaply; the 128 GiB `r5a.4xlarge` EC2 instance we run Plover on currently costs about $650 per month to run continuously, whereas a 32 GiB version of the same instance – which Plater should be able to run comfortably on – is only about $160 per month. Based on our load testing findings, however, multiple Plater instances would likely be required to reach a throughput equivalent to a single Plover instance, meaning that Plater is not necessarily cheaper overall if high concurrent load must be handled.

### Other limitations and areas for future work

Our amortized analysis of query speed was centered around only one query (submitted using different batch sizes); while its results seem to align well with the findings of our other analyses, including additional queries might strengthen this analysis. On a separate note, it is possible that the AWS EC2 instance type we used to host the platforms – which is a “memory optimized” instance type – might be better suited to the Plover platform, given its memory-intensive nature. Hosting Plater on a different kind of instance might be an interesting experiment, though the difference between the two tools in our analysis was large enough that we imagine this would not significantly change the results. Finally, it may be interesting to compare the performance of Plater when deployed with Automat (github:RENCI-AUTOMAT/Automat-server), which distributes queries over multiple Plater instances and thus should be able to handle higher load than a single standalone Plater instance. We suspect that a few Plater instances would be required to – in combination – handle as high a load as a single Plover instance, possibly making Plover more cost-effective for high load scenarios in the end.

## Footnotes

1 Our primary motivation for moving on from Neo4j was performance, in particular query speed; while semantic reasoning can be encoded in Neo4j queries, it tended to cause a notable performance drop in our experience. Convenience of deployment was another factor.

2 Reasoning systems answer more complex queries by stitching together one-hop KG queries.

3 Black text represents specific “pinned” concepts, green text represents edge type constraints, gold text represents node type constraints, purple text represents edge attribute constraints, and asterisks represent a “wildcard” node/edge type. In reality, nodes are referred to only by standard identifiers, rather than their names (e.g., `NCBIGene:672` for the gene *BRCA1*).

4 We ran Plater from a fork containing these modifications, available in GitHub at github:amykglen/Plater/tree/subclasscypher.

5 Both of these modifications were incorporated into Plater v1.6.0. Note that the number of edges in a TRAPI response may be greater than the number of “results”; Plater does not have a straightforward way of limiting the number of answer edges specifically.

6 This corresponds to RTX-KG2 edges with the `domain_range_exclusion` property set to `True`.

7 Queries were deduplicated by creating hash keys for each query that consisted of a string somewhat similar to a Neo4j Cypher query, in which node IDs were canonicalized and sorted, categories/predicates were sorted, and qualifiers were included as well. The deduplication method is available in the `sample_kg2_queries.py` script in the RTX GitHub repository.

8 These times were those taken by the previous RTX-KG2 API software stack, which was not evaluated in this study.

9 ARAX is in the habit of specifying `is_set=True` on all query nodes when querying RTX-KG2, but that is only because the current RTX-KG2 API can process such queries more quickly, and ARAX only needs the answer knowledge graph, not the TRAPI `results`. No other queriers of RTX-KG2 appear to use `is_set=True`, since knowledge providers like RTX-KG2 only answer one-hop (i.e., single-edge) queries.

10 Technically the four subsets of queries were always run in the same order (ITRB Prod, Any, Long, Hand-crafted), but within those subsets queries were run in random order.

11 While this represents a fairly extreme load for a single instance, our RTX-KG2 APIs do occasionally face such high load in practice when being used intensively by, for instance, Every Cure (everycure.org).

12 The `neo4j` directory on the hosting instance is mounted as a volume for the Neo4j docker container.

13 Answer sets were considered identical if they returned the same *edges*; the number of TRAPI *results* differed somewhat, as the two platforms have slightly different (but both valid) ways of constructing results.

14 Coefficient of variation was calculated as *standard deviation/mean* duration for each query.

15 For this query, Plater returned 5,161 nodes, 22,618 edges and Plover returned 5,163 nodes, 22,629 edges.

16 We define “at rest” as when the platform is sitting idle without receiving any queries; we measured this after the build process completed and associated memory had been cleared, but before any queries were sent to the platform.

17 All of the 75 test queries for which Plater returned a 200 status code are included here.

